# The Digital 3D-Atlas MAKER (DAMAKER): a dynamic and expandable digital 3D-tool for monitoring the temporal changes in tissue growth during hindbrain morphogenesis

**DOI:** 10.1101/2021.03.29.437592

**Authors:** Matthias Blanc, Frederic Udina, Cristina Pujades

**Author notes:** corresponding author, Correspondence to: Cristina Pujades, PhD, Department of Medicine and Life Sciences, Universitat Pompeu Fabra PRBB, Dr Aiguader 88, 08003 Barcelona, SPAIN.

## Abstract

Reconstruction of prototypic three-dimensional (3D) atlases at the scale of whole tissues or organs requires specific methods to be developed. We have established a digital 3D-atlas maker (DAMAKER) and built a digital 3D-temporal atlas to monitor the changes in the growth of the tissue and neuronal differentiation domain in the zebrafish hindbrain. DAMAKER integrates spatial and temporal data from cell populations, neuronal differentiation and brain morphogenesis, through *in vivo* imaging techniques paired with image analyses and segmentation tools. First, we generated a 3D-reference from several imaged hindbrains and segmented them using a trainable tool; these were aligned using rigid registration, revealing distribution of neuronal differentiation growth patterns along the axes. Second, we quantified the dynamic growth of the neuronal differentiation domain vs. the progenitor domain, and by *in vivo* neuronal birthdating experiments we generated a digital 3D-temporal map of the neuronal growth in the whole hindbrain, revealing the spatiotemporal dynamics of neuronal differentiation upon morphogenesis. Last, we applied it to glutamatergic and GABAergic neurons, as proof-of-concept that the digital 3D-temporal map could be used as a proxy to infer neuronal birthdate. As this protocol uses open-access tools and algorithms, it can be shared for standardized, accessible, tissue-wide cell population atlas construction.

## INTRODUCTION

Generating a precise temporal cartography of cell populations in the three-dimensional (3D) embryonic brain is essential for understanding how brain morphogenesis impacts the position, and therefore function, of neuronal progenitors and neuronal clusters in normal and pathological conditions. Pioneer work in Drosophila provided 3D-registration atlases and software that can automatically find the corresponding landmarks in a subject brain, and map them to the coordinate system of a target brain (Heckscher et al., 2014; Peng et al., 2011). However, up to now, the incorporation of time —and therefore information of tissue morphogenesis— in 3D-atlases is still one of the main challenges that needs to be solved.

Digital 3D-atlases enable the exploration of brain structures in the 3D-context and can be updated to incorporate new information, which is essential for the exponential increase of knowledge. Existing gene expression 3D-atlases in larval and adult fish provide valuable tools (Chow et al., 2020; Jaggard et al., 2020; Kenney et al., 2021; Ronneberger et al., 2012; Tabor et al., 2019). However, they lack the temporal component, most probably because whole brain images are particularly hard to register across space and time, even though there are recent efforts aiming to make this more accessible (Dsilva et al., 2015; Fernandez and Moisy, 2020). Temporal digital 3D-atlases are essential to understand how the generation of brain cell diversity occurs concomitantly with brain morphogenesis, which results in a dramatic transformation from a simple tubular structure (e.g., the neural tube) to a highly convoluted structure (e.g., the brain). Thus, our goal was to provide an atlas allowing: i) the incorporation of time to enable morphogenesis studies; ii) the quantification of 3D-patterns with a standardized method; and iii) an easy access to the atlas data with user-friendly upgradability, without the need to send the data to the atlas-makers for processing. Such features of a digital atlas are crucial for morphogenetic studies that rely on 3D-topologies to understand how specific regions develop; moreover, to do this in zebrafish would enhance the use of zebrafish as human avatars in neurodevelopmental disorders by enabling to understand how specific territories respond to insults or gene disruptions.

Here, we developed a new “Digital Atlas-MAKER” pipeline (DAMAKER) that fulfills the previous requirements and can be used to create an expandable atlas of virtually any tissue in any organism. Further, it is able to perform data analysis and quantification on a single platform, FIJI, which is already established as the standard for image analysis in the field. The expandable zebrafish digital 3D-atlas was created by combining *in vivo* imaging of distinct transgenic embryos with the Fijiyama registration tool (Fernandez and Moisy, 2020), which allows multi-modal rigid registration, and the Trainable Weka Segmentation (TWS) (Arganda-Carreras et al., 2017) that leverages a limited number of manual annotations and permits data to be automatically segmented after a short training, in order to quantify tissue volumes. We applied DAMAKER to the monitoring of the growth of the neuronal differentiation domain in the context of the whole zebrafish hindbrain —the embryonic brainstem—, to clarify how cell differentiation and morphogenesis might be intertwined. We made use of the embryonic hindbrain, which is the most conserved vertebrate brain vesicle (Murakami et al., 2005) and undergoes tissue segmentation along the anteroposterior (AP) axis leading to the formation of seven rhombomeres (Fraser et al., 1990; Jimenez-Guri et al., 2010; Kiecker and Lumsden, 2005; Krumlauf and Wilkinson, 2021; Pujades, 2020). This morphogenetic process is accompanied with the initial neuronal differentiation, which starts prior 24 hours post-fertilization (hpf) with a pronounced increase from 30 hpf onwards (Voltes et al., 2019). Neuronal differentiation involves changes in cell proliferative capacity and an extraordinary displacement of neurons from their birth site, and occurs while other dramatic changes take place, such as the generation of the brain ventricle (Belzunce et al., 2020; Gutzman et al., 2008; Hevia et al., 2021; Lyons et al., 2003). Although previous work evoked the importance of the order of neuronal differentiation in ascribing final neuronal position within specific circuits (Kinkhabwala et al., 2011; Pujala and Koyama, 2019; Wan et al., 2019), little is known about the role of the neuronal birthdate in the final organization of differentiated neurons within the hindbrain.

Here, we registered experimental whole zebrafish hindbrain images and mapped neurogenesis and neuronal differentiation patterns with a unified spatial representation. We used a confocal microscopy imaging setup and cross-platform registration and segmentation tools. We build up a digital 3D-hindbrain atlas in which signals corresponding to specific territories, neural progenitor cells, neuronal committed precursors, and differentiated neurons were merged. With these combinations, we could associate specific neuronal populations with anatomical landmarks and quantify complex volumetric data. Next, we performed a tissue wide neuronal birthdate analysis by using a method based in photoconverted fluorescent protein tracing *in vivo* (Caron et al., 2008), which permitted to distinguish early-born from late-born neurons in live embryos. This allowed us to develop a dynamic neuronal differentiation map —the expandable temporal digital 3D-atlas— that revealed the remodeling of the neuronal progenitor and neuronal cluster domains over time. By generating these maps, we unveiled that early-born neurons were always located in the inner part of the differentiation domain and surrounded by younger neurons, generating an inner–outer gradient of early-born vs. late-born neurons. Finally, as a proof-of-concept, we demonstrated that this temporal 3D-atlas could be used as a proxy to infer the birthdate of given neuronal populations. We believe that this tool —the digital 3D-atlas maker, DAMAKER— will allow the integration of the information generated in other laboratories and will help ascribing neuronal birthdate upon gene disruption in zebrafish avatars for human diseases. This collaborative integration is essential for expanding our overall knowledge about how the brain is functionally organized, and therefore advancing brain research, medicine and brain-inspired information technology.

## RESULTS

Our scientific interest was to monitor the dynamic growth of the neuronal differentiation domain in the context of the whole zebrafish hindbrain —the embryonic brainstem—, with the emphasis in how neuronal birthdate prefigures position within the differentiation domain.

### Manual analysis revealed the dramatic growth of the hindbrain neuronal differentiation domain at the expense of the progenitor domain

First, we monitored the growth of the neuronal differentiation domain by spatiotemporal analysis. We made use of Tg[βactin:HRAS-EGFP;HuC:Kaede^Red^] embryos, in which the whole tissue cell membranes were labelled in green, and the neuronal differentiation domain with Kaede^Red^. Embryos (n = 5 embryos/stage) were imaged during a temporal interval that encompassed neurogenesis, neuronal differentiation and hindbrain morphogenesis (24—72 hpf), and the size of both the progenitor domain (PD) and the neuronal differentiation domain (NDD) were assessed (Figure S1). The NDD dramatically increased during this time period (Figure S1A—S1D, S1A’—S1D’, S1A’’—S1D’’; see the PD in green and the NDD in magenta), consistent with previous qualitative descriptions (Belzunce et al., 2020; Lyons et al., 2003; Voltes et al., 2019). To assess the volumetric changes, we manually segmented masks and quantified them using a homemade macro, which implied interpolating a dozen of manually segmented sections across the stack (see Supplementary Material 1). Upon volume quantification, we observed that little number of cells underwent differentiation before 24 hpf (V_NDD_ = 1.67 [SD ± 0.26] × 10^6^ μm^3^ vs. V_PD_ = 4.72 [SD ± 0.21] × 10^6^ μm^3^; Figure S1A’— S1A’’). However, this situation dramatically changed few hours later because by 36 hpf the NDD was higher than the PD (V_NDD_ = 6.77 [SD ± 1.08] × 10^6^ μm^3^ vs. 3.45 [SD ± 0.43] × 10^6^ μm^3^; Figure S1B’—S1B’’). By 48 hpf the differentiated neurons occupied mostly the whole hindbrain (V_NDD_ = 10.36 [SD ± 0.72] × 10^6^ μm^3^ vs. V_PD_ = 1.96 [SD ± 0.52] × 10^6^ μm^3^; Figure S1C’—S1C’’), and by 72 hpf the PD was almost entirely depleted (V_NDD_ = 14.48 [SD ± 0.69] × 10^6^ μm^3^ vs. V_PD_ = 0.99 [SD ± 0.32] × 10^6^ μm^3^; Figure S1D’—S1D’’). Thus, differentiated neurons populated the whole hindbrain in approximately 48 hours. This remodeling and drastic shift in the balance between progenitor and differentiated neuronal populations could be better observed by plotting the ratios of the PD and the NDD over time (Figure S1E). Although this manual analysis was useful in this specific case because the differences in size of the PD and NDD were important, more subtle changes could not be assessed. Thus, we confirmed that if we wanted to transition towards quantitative biology, we needed a more elaborated pipeline, which diminished the weaknesses associated with manual segmentation and the lack of reliability of spatial features due to interindividual variability. To overcome this, we developed a standardized and reliable 3D-imaging processing method that allowed to generate digital 3D-models from the averaging of multiple samples and to quantify volumes in a reliable manner, the digital 3D-atlas maker or DAMAKER.

### Developing a standardized, reliable 3D-image processing and quantification method

Building upon previous work that reconstructed prototype 3D-atlases of whole embryos in Drosophila (Heckscher et al., 2014; Peng et al., 2011) and of larval or adult zebrafish (Chow et al., 2020; Jaggard et al., 2020; Kenney et al., 2021; Ronneberger et al., 2012; Tabor et al., 2019), we aimed to create a digital 3D-atlas of the hindbrain neuronal differentiation domain that incorporated time. To build it up we designed a pipeline requiring the following steps: i) registration of *in vivo* 3D-images displaying specific signals to align raw samples from different embryos (Figure 1A), ii) segmentation of the 3D-images to generate labeling masks, each corresponding to a given marker or cell population (Figure 1B), and iii) processing these masks to create average digital 3D-models (Figure 1C). In addition, the pipeline should allow to segment and isolate specific signals from *z*-stack images in a semi-automatic manner in order to i) quantify the desired signal in different embryos (Figure 1D–1E), ii) generate a reference hindbrain digital 3D-model, iii) use this reference 3D-model for further registrations and comparisons (Figure 1F), and iv) combine reference 3D-models to generate a digital 3D-atlas. In our case, to establish a reference tissue signal we made use of Tg[HuC:GFP] embryos, in which GFP is uniformly displayed in the whole neuronal differentiation domain, helping to reference its morphology. Then, several signals could be used, such as those labeling specific domains, cell populations, etc (Figure 1A). To apply this pipeline, acquired 3D-images from different Tg[HuC:GFP] embryos (e1–eY; *n* = 5 each) were registered to the reference by Fijiyama registration, in which rigid registration was favored, although non-rigid registration could be considered at the cost of sample shape variability. Once the alignment of the HuC signal was accomplished, the obtained transformation matrix could be applied to other signals (neurog1, gad1b, …) (Figure 1A). After alignment of 3D-images, tissue segmentation was applied in order to generate binary masks for each labeling/staining (Figure 1B). Here, segmentation settings were at tissue scale, both in an effort to keep compute power low for pipeline accessibility and for our interest in cell populations, although single cell resolution could be achieved with nuclear reporter lines. In order to discern noise vs. signal in the raw data from each embryo, different users (u1–uX; *n* = 5) were invited to employ the Trainable Weka Segmentation (TWS) tool to train classifiers (one per user) (Figure 1B). We used a single classifier to segment all signals, but in the case of low or sparse signals (or specific structures within a signal) separate classifiers would be better for leveraging the structural recognition capabilities of TWS. To reduce the potential impact of the human bias from the trained classifiers, we averaged the resulting segmented images from multiple users’ trained classifiers (Figure 1B). In the case of having an outlier user, this classifier was not considered (as example, see u4 in Figure 1B). Accordingly, we generated as many binary 3D-masks as users (*n* = 4; Figure 1B). The users’ averaged embryo 3D-masks displayed a gradient, which represented the spatial similarity between users’ segmentations. We selected the signals agreed by the four users (Figure 1B), although this could be differently thresholded to display signal consensus among users, with the strength of the threshold setting up the diversity. For image processing, these embryo 3D-masks were averaged to merge information from all embryos (e1–eY; *n* = 5) to obtain the “label 3D-model”. At the end, we generated as many label 3D-models as desired signals, by reiteration of the whole process (Figure 1C). The digital label 3D-model displayed a gradient, which represented the spatial similarity between embryos. To establish the most accurate consensus selection, we compared label 3D-models thresholded for different levels of consensus between embryos. We selected the signals displayed by at least 3 embryos, compared the distribution of the labeling between embryos, and built up the digital reference 3D-atlas. Thus, this pipeline allowed us to: i) assess signal similarity between segmentations to limit the bias due to the individual subjectivity, ii) obtain reliable quantification at the tissue level, iii) produce a representative model of a label, iv) estimate interindividual variability, and v) build up a reference brain to be further used for spatiotemporal comparisons.

**Figure 1:**
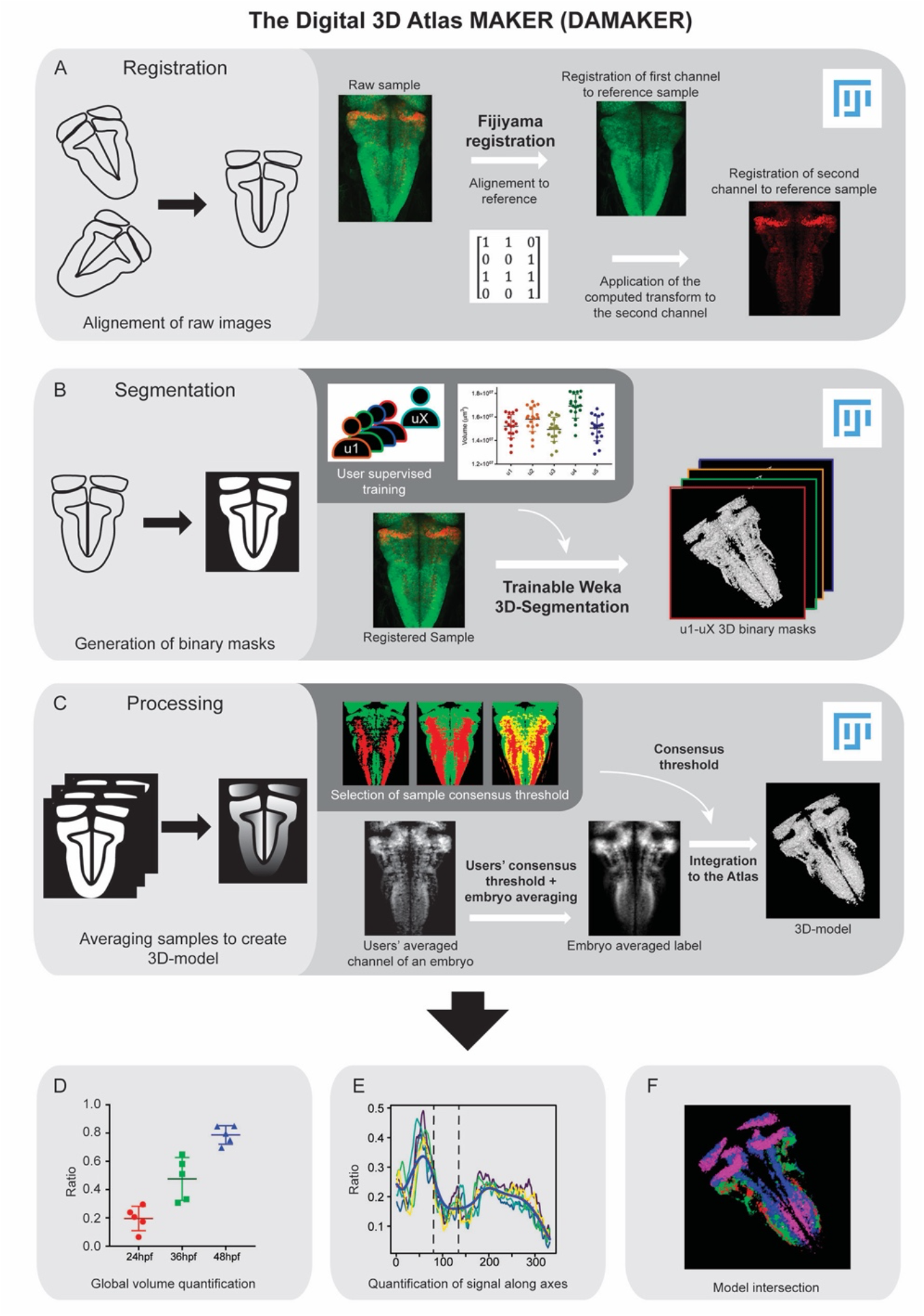
Image analysis pipeline (DAMAKER) and digital 3D-atlas construction. Schematic depiction of sample processing through the digital 3D-atlas making pipeline. (A) For image registration, acquired dual channel images from embryos are registered using the Fijiyama registration plugin in FIJI, the first channel (Huc:GFP signal in green) is aligned to a previously selected HuC signal which serves as reference of orientation. The resulting transform file is then used to align the signal of interest (here, vglut2a:DsRed in red). (B) For image segmentation, Tg[HuC:Kaede] embryos displaying magenta in the early differentiated neurons and green in later differentiated neurons are used for supervised training of the Trainable Weka 3D-Segmentation algorithm. Images are provided to users, and each user brush strokes on areas with signal and areas considered as background across the stack, in order to obtain a classifier (e.g., a set of filters and parameters to be used by the algorithm to isolate the signal). Users’ generated classifiers are compared by quantification of a dozen of HuC-signals from various embryos. User classifiers resulting in more than 10% variability among users (ANOVA test) were deemed to be outlier (u4, user 4) and were not used for segmentation of samples. For each embryo, as many 3D-binary masks as user-generated classifiers were created. (C) For processing, 3D-binary masks of a single embryo are averaged to obtain a users’ average segmentation; this allows to assess the interuser segmentation variability and limits the human bias introduced by the supervised training. Signal similarity threshold between users can then be selected (here, consensus among all users was isolated). Users’ average embryo 3D-models of a given condition (*n* = 5) are then averaged together to obtain the embryo’s averaged label. To select sample consensus threshold, we compared embryo’s averaged labels at different thresholds with their negative imprint onto the neuronal differentiation domain. Signal consensus among at least three embryos seems to recapitulate the best of the original signal, with marginal overlap between the signal and its imprint. Sample consensus threshold is then applied to the embryo’s averaged label to generate the final 3D-model, ready for the integration into the atlas. Several functions can be performed with DAMAKER such as (D) quantification of embryo averaged label volumes in μm3 and displayed them as a ratio of signal over the neuronal differentiation domain; (E) quantification of the signal along axes by virtually reslicing embryo averaged models along the axes and quantifying volumes at each position; (F) intersection of the 3D-model integrated in the atlas to allow qualitative observations of overlapping or segregation of signals.

### Building a multilayered digital 3D-atlas

Next, we wanted to add multiple layers to the 3D-atlas of the neuronal population domain built at 72hpf, the stage in which most of the cells have undergone neuronal differentiation. For this, we made use of double transgenic embryos displaying a fluorescence reporter in the neuronal differentiation domain, used as a reference for alignment in one channel, and another fluorescence reporter in the specific cell population or territory of interest (Figure 2). We first incorporated territory landmarks to generate a spatial coordinate system on which to have a quantitative representation of the given cell population distribution across the tissue. At 72 hpf, we imaged Tg[Mu4127;HuC:GFP] embryos, which displayed DsRed in discrete hindbrain domains such as rhombomeres (r) 3 and 5 (Distel et al., 2009), and GFP in the neuronal differentiation domain. Thus, we could have the r3 and r5 anatomical landmarks along the anteroposterior axis within the neuronal differentiation domain (Figure 2A–2A’’). When rendered, we could display the 3D-view of the total r3 and r5 volumes (Video 1A). To monitor possible differences in the spatial distribution within the neuronal differentiation territory, we quantified the signal volume across body axes using this pipeline. For this, we took sample segmentations and created virtual “slices” along the different axes and quantified the signal ratio at each “slice” (Figure 2B–2B’; the number of virtual slices along the given body axis is indicated on the X-axis). We observed that the signal distribution was concentrated in r3 and r5 along the anteroposterior (AP) axis (Figure 2B), whereas it was similarly distributed along the mediolateral (ML) axis (Figure 2B’).

**Figure 2:**
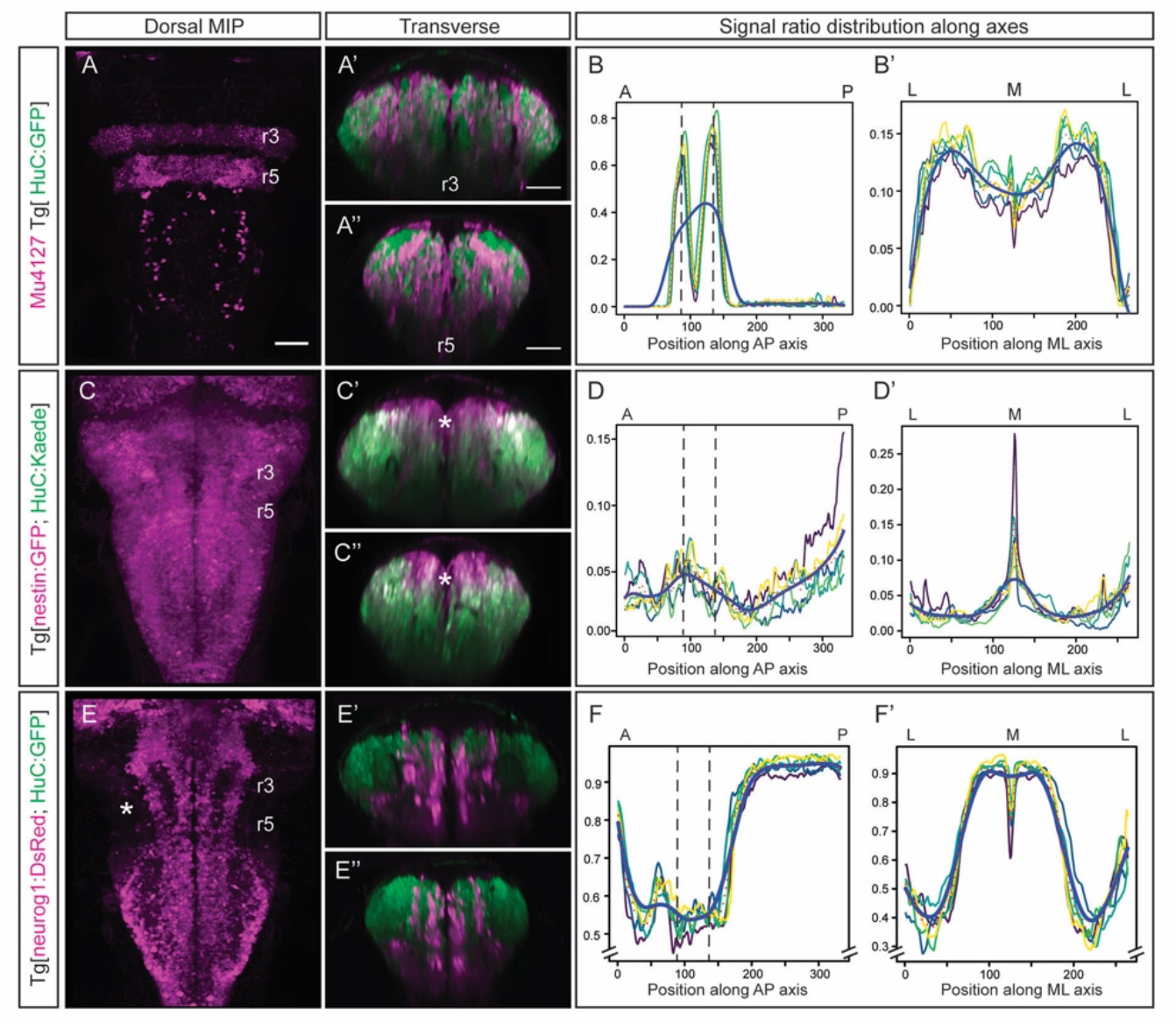
Incorporating territory landmarks allows precise spatial characterization of signals. (A–A”) Dorsal maximal intensity projection (MIP) and transverse views through r3 and r5 of registered Mu4127 Tg[HuC:GFP] embryos imaged at 72hpf. (B–B’) Signal ratio distribution along the anteroposterior (AP) and mediolateral (ML) axes, respectively. (C–C”) Dorsal MIP and transverse views through r3 and r5 of registered Tg[nestin:GFP;HuC:Kaede] embryos imaged at 72hpf. Note the white asterisk indicates the small territory nestin-positive and HuC-negative, which corresponds to the progenitor domain almost depleted at this stage. Nestin-positive domain depicts the neural progenitor derivatives. (D–D’) Signal ratio distribution of progenitor nestin signal along the AP and ML axes, respectively. (E–E”) Dorsal MIP and transverse views through r3 and r5 of registered Tg[neurog1:DsRed;HuC:GFP] embryos imaged at 72hpf. Note the white asterisk in the dorsal MIP views indicates the anterior hindbrain, which displays lower neurog1 signal. (F–F’) Signal ratio distribution of neurog1 signal along the AP and ML axes, respectively. Note the offset in the Y-axis for easier readability of the signal variations. Dorsal MIP display anterior to the top. Scale bar, 50μm. In (B–B’, D–D’, F–F’) X-axis displays the number and position of virtual slices along the AP (B, D, F) and ML (B’, D’, F’) axes. Colored lines correspond to different individual embryos (*n* = 5), red dotted line to the embryos’ average, and solid blue line to the non-linear regression. Black dashed lines parallel to the Y-axis correspond to r3 and r5 positions revealed by Mu4127 signal.

For assessing the size of the progenitor cell domain we used Tg[nestin:GFP;HuC:KaedeRed] embryos, in which the territory nestin-positive and HuC-negative corresponded to the progenitor domain (Figure 2C’–2C’’, see white asterisk). This allowed us to generate the 3D-model of the progenitor domain demonstrating that by 72 hpf this domain was extremely reduced (Video 1B). When volumes were automatically quantified and the ratios along axes plotted, we observed that the progenitor domain was mostly depleted at 72 hpf (Figure 2D– 2D’), similar to the previously observed results (Figure S1). Next, we investigated the distribution of neuronal committed progenitors, according to their proneural gene expression (Guillemot, 2007), in this case *neurog1*. In Tg[neurog1:DsRed;HuC:GFP] embryos, we observed the striped-pattern distribution of *neurog1-*progenitor derivatives in the neuronal differentiated domain (Figure 2E–2E’’; Video 1C). When automatic quantification of the *neurog1*-volume was acquired, we observed a change in the ratio of labeling along the AP axis (Figure 2F), supporting the observation of regions with less *neurog1*-derivatives in the anterior hindbrain when compared with the spinal cord (Figure 2E, see white asterisk). In addition, the more medial distribution of *neurog1*-derivatives was shown (Figure 2F’). Thus, this quantitative analysis allowed to associate signal distribution to specific territories or cell populations.

Next, we wanted to map distinct neuronal differentiation programs. First, we analyzed the spatial distribution of motoneurons by using Tg[isl1:GFP;HuC:Kaede^Red^] embryos and observed their ventral position as previously described (Figure 3A–3A’’; Video 2A). Upon quantification, higher signal was observed in discrete patterns along the AP axis with a clear increase in the spinal cord (Figure 3B) due to an earlier onset, and the accumulation of motoneurons in the more medial domain (Figure 3B’). Then, we explored the specific neurotransmitter-expressing cell populations, considering the mainly expressed markers of excitatory (glutamatergic neurons) and inhibitory populations (GABAergic neurons) in the zebrafish hindbrain (Higashijima et al., 2004). Using Tg[vglut2a:DsRed;HuC:GFP] and Tg[gad1b:DsRed;HuC:GFP] embryos we could observe the main contribution of glutamatergic neurons to the most dorsolateral differentiation domain of the hindbrain (Figure 3C–3C’’), whereas GABAergic neurons allocated more medially (Figure 3E–3E’’). Upon building the digital 3D-maps (Videos 2B–2C), we quantified the distribution along the axes and unveiled a smaller contribution of glutamatergic neurons to the r3-r5 region (Figure 3D), whereas GABAergic signal was enriched in r2 (Figure 3F) when compared to the rest of the hindbrain at this stage. This quantification prettily recapitulated the different mediolateral distribution of these two neuronal populations (Figure 3D’, 3F’), as previously established as spatially restricted domains of neurotransmitter expression (Kinkhabwala et al., 2011; Koyama et al., 2011; Pujala and Koyama, 2019). In addition, the overlay of the distinct digital 3D-maps provided information about the segregation of populations (Video 2D). Overall, the analysis of reporter fluorescent proteins expression allows the virtual colocalization of signals, and therefore the model intersection in order to study specific subsets of neuronal populations.

**Figure 3:**
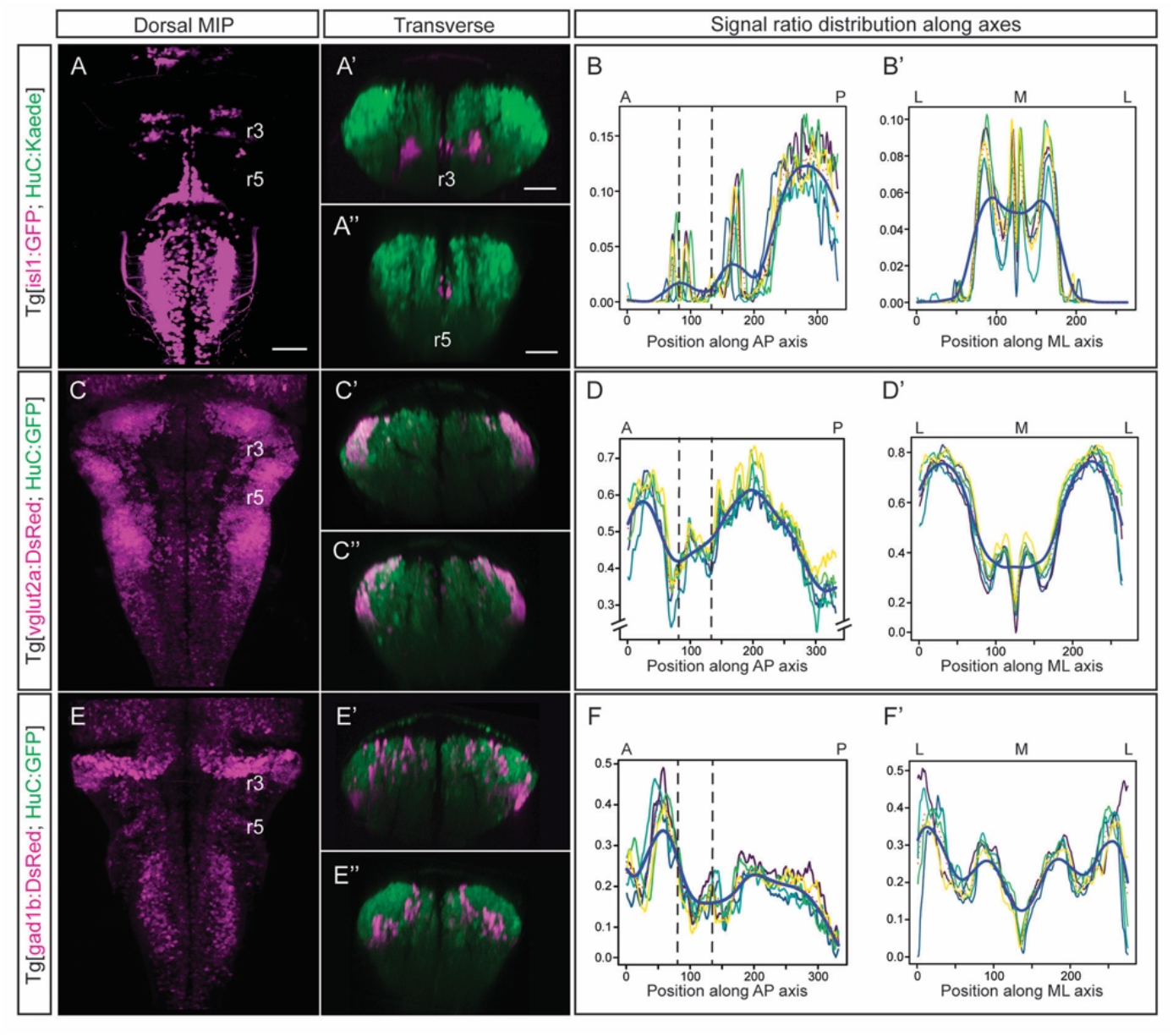
Quantification of the signal along axes foreshadows the distinct neuronal cell populations distribution. (A–A”) Dorsal MIP with contrast enhancement to make r3 signal more visible, and transverse views through r3 and r5 of registered Tg[isl1:GFP;HuC:Kaede] embryos imaged at 72hpf. Note the ventral position of motoneurons. (B–B’) Signal ratio distribution of isl1 along the AP and ML axes, respectively. (C–C”) Dorsal MIP and transverse views through r3 and r5 of registered Tg[vglut2a:DsRed;HuC:GFP] embryos imaged at 72hpf. (D–D’) Signal ratio distribution of vglut2a along the AP and ML axes, respectively. (E-E”) Dorsal MIP and transverse views through r3 and r5 of registered Tg[gad1b:DsRed;HuC:GFP] embryos imaged at 72hpf. (F–F’) Signal ratio distribution of the gad1b signal along the AP and ML axes, respectively. Dorsal MIP display anterior to the top. Scale bar, 50μm. In (B–B’, D–D’, F–F’) X-axis displays the number and position of virtual slices along the AP (B, D, F) and ML (B’, D’, F’) axes. Colored lines correspond to different individual embryos (*n* = 5), red dotted line to the embryos’ average, and solid blue line to the non-linear regression. Black dashed lines parallel to the Y-axis correspond to r3 and r5 positions revealed by Mu4127 signal distribution.

### Building a temporal digital 3D-atlas of neuronal differentiation

Previous work demonstrated the importance of neuronal birthdate in ascribing neuronal function: i) the stripe patterning of *alx*-expressing neurons, which are involved in swimming behavior, is age-related (Kinkhabwala et al., 2011), and ii) the position of the V2a cell body and the order of spinal projections in the hindbrain relies on their neuronal birth (Pujala and Koyama, 2019). Thus, we wanted to explore whether neuronal birthdate was a general rule that dictated the spatial distribution of differentiated neurons within the hindbrain.

We performed spatiotemporal analyses of neuron birthdating using Kaede photoconversion experiments as a tool to register temporality (Caron et al., 2008). In this case, Kaede^Green^ in Tg[HuC:Kaede] embryos was *in vivo* photoconverted to Kaede^Red^ at different developmental stages (24, 36 or 48hpf), and embryos were imaged at 72 hpf when the majority of the progenitor cells were extinguished (Figure 4A–4C, 4A’–4C’, 4A’’–4C’’). This allowed us to assess the relative position of early-born neurons (Figure 4; Kaede^Red^-cells in magenta) vs. late-born neurons (Figure 4; Kaede^Green^-cells in green). We observed that early differentiated neurons were more ventrally and medially allocated than late differentiating neurons, which piled up (Figure 4A—4A’’, 4B—4B’’, 4C—4C’’; see magenta vs. green cells in transverse sections). To determine whether the temporal order of neuronal differentiation prefigured the spatial distribution of neurons in the tissue, we compared the label 3D-models of the neuronal domains generated before 24 hpf, 36 hpf and 48 hpf in the context of the whole differentiated neuronal domain at 72 hpf (Videos 3A–3C; see youngest neurons in yellow, the second-youngest neurons in blue, the bit older neurons in green). We did overlap and color-coded them according to the neuronal birthdate (Figure 4D—4D’’’; see youngest neurons in yellow, neurons differentiated between 36 and 48 hpf in blue, neurons differentiated between 24 and 36hpf in green, and the oldest ones in magenta; note that magenta neurons were already differentiated at 24 hpf). We observed that between 24 and 36 hpf, cells differentiated all along the AP axis (Figure 4B–4B’’, 4D–4D’’; see green domain), and that an important part of neurons born between 48 and 72 hpf were located in the most anterior domain of the hindbrain (Figure 4D). This differentiation capacity diminished with time (Figure 4D; see small blue domain); indeed, by 48 hpf, almost the whole differentiation domain was already occupied by previously-differentiated neurons (Figure 4C’–4C’’, 4D’–4D’’’).

**Figure 4:**
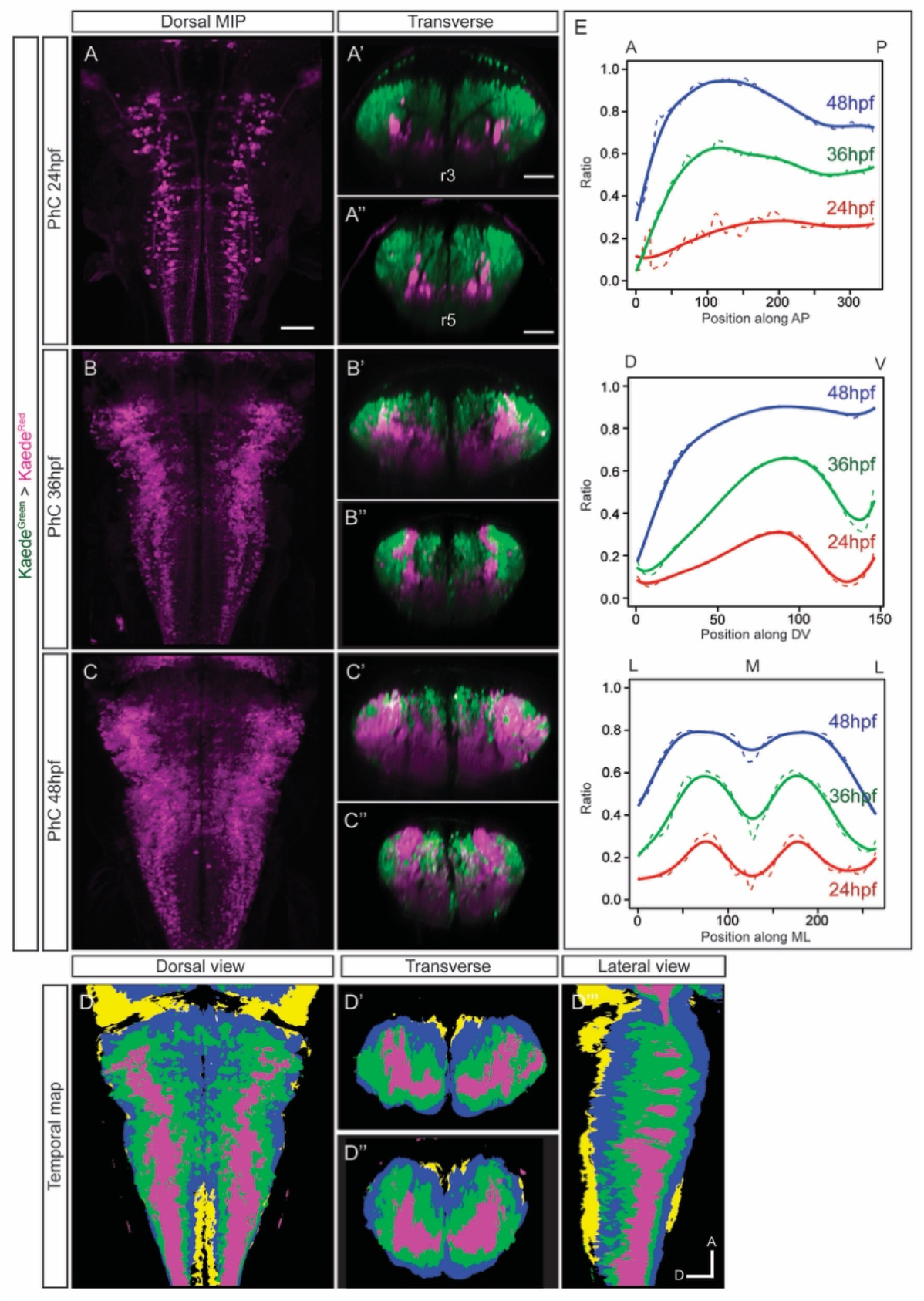
Differentiated neurons spatially organize according to their birthdate. (A–C) Dorsal maximal intensity projection (MIP), and (A’–C’, A’’–C’’) transverse views through r3 and r5 of registered Tg[HuC:Kaede] zebrafish embryos acquired at 72hpf after photoconversion of Kaede^Green^ at the indicated times (A–A’’, 24 hpf; B–B’’, 36 hpf; C–C’’, 48 hpf). Note that Kaede^Red^ is displayed in neurons born before the time of photoconversion, and Kaede^Green^ in neurons born after each photoconversion time. (D–D’’’) Dorsal, transverse and lateral views of the temporal map resulting from the intersection of photoconversion 3D-models. Neurons differentiated before 24hpf are displayed in magenta, neurons differentiated between 24 and 36hpf are colored in green, neurons differentiated between 36 and 48hpf are in blue, and those differentiated between 48 and 72hpf in yellow. A, anterior; D, dorsal. Dorsal MIP display anterior to the top. Scale bar, 50μm. (E) Volumes of late-born and early-born neurons were quantified along the AP, dorsoventral (DV) and ML axes. X-axis displays the number and position of virtual slices along the given body axis. The ratio of early-born neurons over the total neuronal differentiation domain was plotted as mean (segmented) and non-linear regression (solid) (*n* = 5 embryos). Color-coded lines correspond to the indicated embryonic stages.

The overlaying maps showed that i) most of the neuronal differentiation events occurred between 24 and 48 hpf, and ii) early-born neurons were always located in the inner part of the differentiation domain and surrounded by younger neurons, generating an inner–outer gradient of early-born vs. late-born neurons. This suggested that the position of neurons relied on their neuronal birthdate, and demonstrated the importance of morphogenesis for remodeling the differentiation domain. When the volumes of neurons born at different times were quantified at 72 hpf, we observed that only a small part of the NDD came from neurons born before 24 hpf (3.63 [SD ± 1.30] × 10^6^ μm^3^ for early-born neurons vs. 12.81 [SD ± 2.15] × 10^6^ μm^3^ for late-born neurons). Most neurons underwent differentiation between 24 hpf and 48 hpf, with half of the differentiated neurons arising before 36 hpf (7.8 [SD ± 1.7] × 10^6^ μm^3^ neurons born before 36 hpf vs. 7.08 [SD ± 1.43] × 10^6^ μm^3^ after 36 hpf) and the rest were differentiated between 36 and 48 hpf (11.65 [SD ± 1.42] × 10^6^ μm^3^ neurons born before 48 hpf vs. 3.19 [SD ± 0.43] × 10^6^ μm^3^ after 48 hpf). This continuous birth of differentiated neurons could be better observed by assessing the relative contribution of early-vs. late-born neurons as compared to the whole NDD. When the ratio of the NDD over the whole hindbrain was assessed, a steady increase of neuronal differentiation was observed (24 hpf: 0.20 ± 0.09; 36hpf: 0.48 ± 0.15; 48 hpf: 0.79 ± 0.07). Overall, these results indicated a consistency of birthdate patterns across samples, suggesting that the positions of differentiated neurons in the inner–outer differentiation gradient was linked to their birthdate.

To monitor possible differences in the spatiotemporal distribution within the neuronal differentiation domain, we took sample segmentations from the birthdate experiments (Figure 4A—4C), and created virtual “slices” along the different axes to sample and quantify the signal throughout the hindbrain (Figure 4E; the number of virtual slices along the given body axis is indicated on the X-axis). We observed that the domains corresponding to early-vs. late-born neurons followed the same trend (e.g., the red, green and blue lines did not display major changes in the multiple virtual slices), with no main differences along the AP or ML axes (Figure 4E). The abundance of youngest neurons in the most anterior parts of the hindbrain was confirmed by the AP distribution (Figure 4E, see the high ratio in the most anterior domain at 48 hpf). As expected, when the analysis was performed along the DV axis of the neuronal differentiation domain, differences in cell allocation relying on the neuronal birthdate could be appreciated, with early-born neurons (24 hpf) displaying a ventral enrichment (Figure 4E, see DV-axis graph). Overall, these whole population analyses allowed us to better understand the intertwinement of morphogenesis and cell behaviors. Specifically, the combination of *in vivo* imaging techniques with image analysis tools to construct the digital 3D-models revealed the remodeling of the neuronal progenitor and neuronal cluster domains upon time.

### The digital 3D-atlas of temporal differentiation as a proxy to infer neuronal birthdates

Our next goal was to investigate whether position in the neuronal differentiation domain at late embryonic stages could be used as a proxy for neuronal birthdating. As a proof-of-concept, we used the Tg[vglut:DsRed] and Tg[gad1b:DsRed] lines, which display DsRed in the glutamatergic and GABAergic neurons, respectively (Satou et al., 2013). To dissect when glutamatergic neurons were born, we generated a label 3D-model of Tg[vglut2a:DsRed] embryos at 72 hpf, and overlaid it on the temporal digital 3D-maps created in Figure 4, in such a manner that glutamatergic neurons were color-coded according to their birthdate (Figure 5A—5A’’). Glutamatergic neurons highly contributed to the differentiation domain, and were born during all the analyzed temporal windows (Figure 5A–5A’’; Video 4A). They displayed the inner-outer distribution depending on birth-time, with blue cells in the outer part and red cells in the inner domain (Figure 5A’–5A’’; Video 4A). These observations suggested that an important part of glutamatergic neurons was born between 24 and 36 hpf (Figure 5A–5A’’; Video 4A; see green cells); moreover, the results indicated that some cells were born before 24 hpf (Figure 5B–5B’’; Video 4B; see red cells). In the case of GABAergic neurons, we took the same approach and overlaid the 3D-model of Tg[gad1b:DsRed] embryos on the temporal digital 3D-map, and color-coded them according to the differentiation time. The neuronal differentiation territory allocated GABAergic neurons born before 48 hpf (Figure 5B–5B’’; Video 4B; see red, green and blue cells), with most neurons born between 24 and 36 hpf (Figure 5B–5B’’; Video 4B; see green cells). In addition, some GABAergic neurons in r3 and r5 were born as early as 24 hpf (Figure 5B–5B’’; Video 4B; see red cells). Overall, this analysis indicated that position of differentiated neurons at 72 hpf could recapitulate neuronal birthdates. To demonstrate that indeed this could be used as a proxy to infer birthdate, and that glutamatergic and GABAergic neurons were born so early, we performed *in situ* hybridizations with *vglut2a* and *gad1b* probes at early stages of embryonic development. We observed that, indeed, glutamatergic and GABAergic neurons were already specified at 24 hpf (Figure 5C, 5D–5D’), although the transgenic lines did not display the reporter expression at this stage. Thus, these results demonstrate that the temporal differentiation 3D-map can be used as a proxy to infer neuronal birthdate, and widens the use of the temporal digital 3D-atlas.

**Figure 5:**
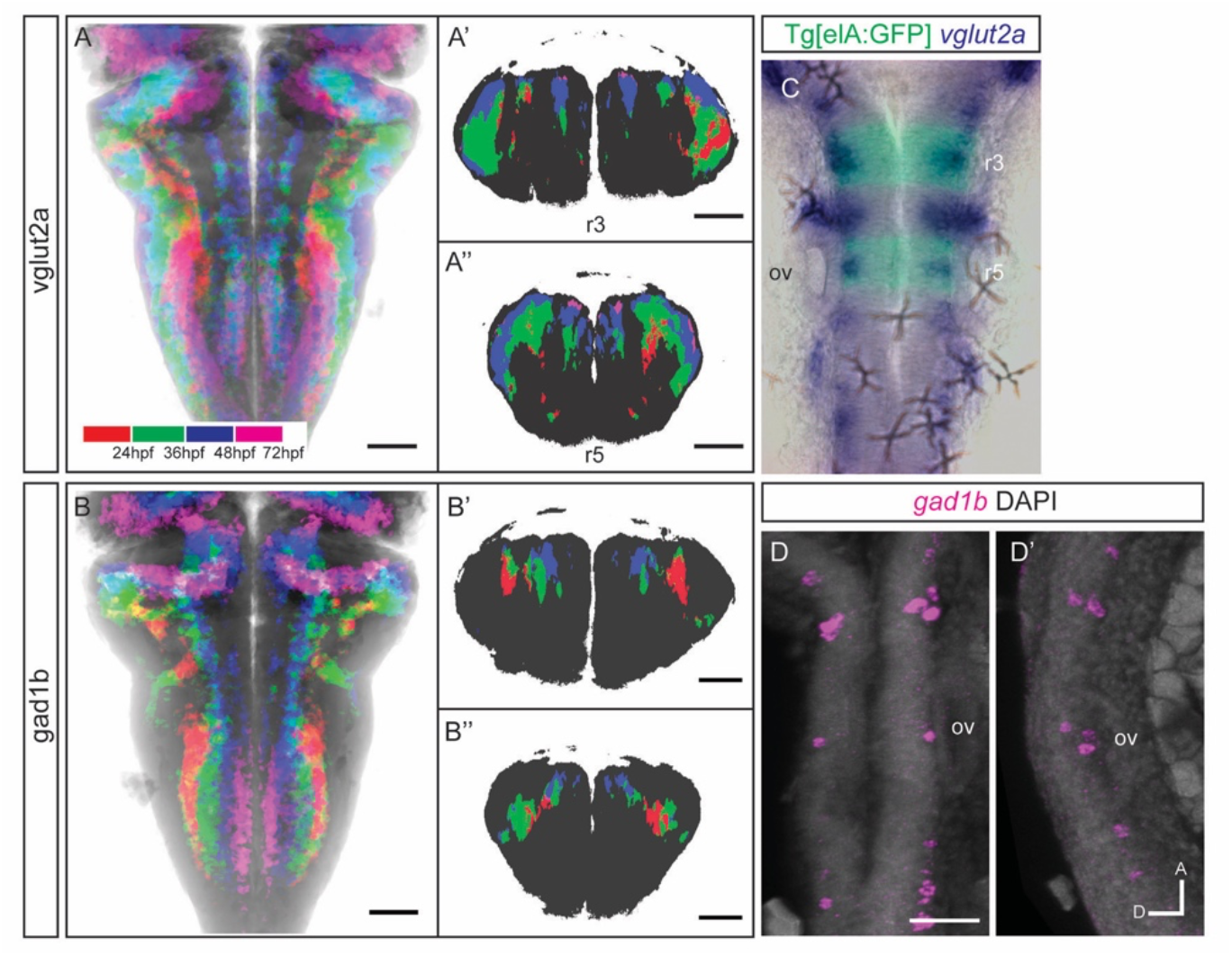
The temporal registration 3D-atlas can be used as a proxy to reveal the birthdate of the desired neuronal differentiated neurons. (A–B) Dorsal average intensity projection (AIP) and (A’–A’’, B’–B’’) transverse views through r3 and r5 of the glutamatergic and GABAergic 3D-models intersected with the temporal 3D-map. vglut2a or gad1b neurons produced before 24 hpf, between 24 and 36hpf, between 36 and 48hpf and between 48 and 72hpf are depicted in red, green, blue and magenta, respectively. The differentiated domain is depicted in black. (C–D) Dorsal MIP of *vglut2a in situ* hybridization of a 24hpf Tg[elA:GFP] embryo, displaying GFP in r3 and r5. (D) Dorsal and lateral MIP of *gad1b in situ* hybridization of 24 hpf embryo. Scale bar, 50μm. ov, otic vesicle. Dorsal and lateral views display anterior to the top.

## DISCUSSION

One of the main current challenges in biology is the 3D-anatomical reconstruction of the tissues’ growth over time and the reconstruction of cell lineage histories. This requires to extract meaningful biological information from large imaging datasets, and some of the bottlenecks are i) the amount of required computing power for the visualization and analysis of the data volumes, and ii) the need of user-friendly tools to be paired with imaging techniques. Here, we developed a digital 3D-atlas maker (DAMAKER), which is a dynamic and expandable 3D-tool for monitoring the temporal changes in growth during tissue morphogenesis. We built a digital representation of the developing vertebrate hindbrain from 3D+time imaging data, which can be used for further quantitative analysis and multilevel modeling. Our goal is to share an established protocol using open-access tools and algorithms that allow standardized and accessible tissue-wide cell population atlas construction and can be used to create an expandable atlas of any tissue in any organism.

In this work, our interest was to explore how neuronal differentiation and morphogenesis might be intertwined during hindbrain development. For this, we registered experimental whole hindbrain images and mapped neuronal differentiation patterns with a unified spatial representation, using confocal imaging setup and developing cross-platform segmentation and registration tools. To improve comparison of various signals across embryos that requires an alignment/registration, which is often done manually and is very time-consuming, we used the Fijiyama plugin in FIJI (Schindelin et al., 2012). This allowed a multi-modal automatic rigid registration process and a large range of customization, and it permits the application of a previously computed transform; namely, an alignment can be computed using one channel of an embryo matching the chosen reference and be applied to the other channel which shares no spatial features with the reference signal. This permits the use of virtually any signal previously added to the atlas, as reference to align others. The quantitative evaluation of datasets often involves manual annotation, which is time-consuming, introduces biases and often constitutes the major constraint in an evaluation pipeline. To overcome this problem, we used the Trainable Weka Segmentation 3D (TWS, (Arganda-Carreras et al., 2017), which combines the image processing toolkit FIJI with the state-of-the-art machine learning algorithms provided in the latest version of the data mining and machine learning toolkit Waikato Environment for Knowledge Analysis (WEKA) (Hall et al., 2009). This tool is modular, allows sophisticated data mining processes to be built from the wide collection of base learning algorithms and tools provided, is open-sourced, and can be easily used as a plugin of FIJI. User trained classifiers, which are a set of filters and thresholds determined by the algorithm to best match input from the user, can be used repeatedly to segment each dataset. By keeping our pipeline in the FIJI ecosystem, we aim at providing a user-friendly tool to share data, upgrade methodology and adapt the pipeline to a specific experimental set up (model organism, organ etc), thereby making it accessible to anyone.

The digital 3D-brain atlases are essential for neurobiology because they allow to address new questions that can not be answered by the use of the available 2D-atlases. The addition of time as a key parameter is an asset, since it will permit to follow cells upon time, and therefore cell displacements and tissue growth. Thus, this temporal registration provides important advantages, as it allowed us to: i) dissect the final location of birthdated cell populations, ii) obtain a whole-cell population coverage, and iii) maintain the *in vivo* positioning of cells within the structure. Until now, most of the currently used approaches either lack whole-cell population coverage, have poor temporal precision or are limited by expression timing of the gene or transgene; more importantly, although they permit coverage of the whole cell population, they do not allow the user to foreshadow the birthdate of a given cell population at the time of interest.

Usually, expression of the reporter genes in transgenic lines is delayed as compared to the gene expression onset, unless the reporter has been inserted within the given gene locus. Now, we can specifically knock-in genes within a targeted genome site thanks to the CRISPR-Cas9 technology, but most of the available—and very valuable—transgenic lines were (and still are) generated with random insertion systems that do not always recapitulate the right onset of gene expression. Thus, our tool can be very useful for predicting neuronal birthdates just by analyzing the position of neurons in the differentiation domain. Besides that, this developed digital platform will be useful to interrogate phenotypes in neurological diseases, and zebrafish avatars from human neurological disorders could be analyzed in a very simple manner. This can provide computational models allowing to merge information from several high-throughput experiments to enlarge our reductionist view, and implement these digital tools for neuronal disorder studies.

In addition, brain morphogenesis studies could use our digital atlas-maker tool, which precisely maps cell populations in the embryonic brain over time and can be easily shared. In the future, DAMAKER can be further improved by using batch processing algorithms (e.g., to assess the effects of pharmacological treatments or gene disruptions) or by adjusting for more geometrically complex tissues or for tissues with less alignment features (such as heart). Last but not least, this pipeline could be applied to time-lapse videos by repeating the process for each frame.

In summary, DAMAKER provides an established protocol using open-access tools and algorithms and allows construction of standardized, accessible, tissue-wide cell population digital 3D-atlases. This approach will help us to fill the knowledge gap between gene regulatory networks, cell poputions and tissue architecture, as it will allow to follow the dynamic events that pattern the early embryonic organs as they happen within the intact system.

## MATERIALS AND METHODS

### Zebrafish strains

Zebrafish (*Dario rerio*) were treated according to the Spanish/European regulations for the handling of animals in research. All protocols were approved by the Institutional Animal Care and Use Ethic Committee (Comitè Etica en Experimentació Animal, PRBB) and the Generalitat of Catalonia (Departament de Territori i Sostenibilitat), and they were implemented according to European regulations. Experiments were carried out in accordance with the principles of the 3Rs.

Embryos were obtained by mating of adult fish using standard methods. All zebrafish strains were maintained individually as inbred lines. The Tg[βactin:HRAS-EGFP] line has GFP in the plasma membrane and was used to label the cell contours (Dale and Topczewski, 2011). Mü4127 transgenic line was used for repairing rhombomeres 3 and 5; it is an enhancer trap line in which the trap KalTA4-UAS-mCherry cassette was inserted into the 1.5Kb region downstream of *egr2a/krx20* gene (Distel et al., 2009). The Tg[nestin:EGFP] labels neural progenitors and derivatives (Lam et al., 2009), and Tg[neurog1:DsRed] (Drerup and Nechiporuk, 2013) labels neuronal committed cells and the *neurog1*-derivatives. Tg[Isl1:GFP] line labels motoneurons (Higashijima et al., 2000). The Tg[HuC:GFP] (Park et al., 2000) and Tg[HuC:Kaede] (Harrison et al., 2014) lines were used to label the whole neuronal differentiated domain. To label GABAergic and glutamatergic neurons, Tg[gad1b:lox-DsRed-lox-EGFP] and Tg[vglut2a:lox-DsRed-lox-EGFP] transgenic lines (Satou et al., 2013) (denoted Tg[gadb1:DsRed] and Tg[vglut2a:DsRed] herein) were used, respectively.

### Confocal imaging samples

Zebrafish embryos were treated with PTU at 24 hpf prior to being anesthetized with tricaine and mounted dorsally in 0.9% low melting point agarose on glass-bottom Petri dishes (Mattek) at the desired time. Images were acquired with a Leica SP8 system using PMT detectors and a 20× glycerol immersion objective, HCX PL APO Lambda blue 20×/0.7 argon laser 30%, Diode 405nm (DPSS 561; Ready Lasers). *z*-stacks were recorded with 0.57×0.57×1.19μm voxel size. Gain was adjusted to each signal in order to display as little burnt signal as possible.

Stained fixed samples were imaged on a Leica SP8 inverted confocal microscope with 20× glycerol immersion objective and hybrid detectors. For x*yz* confocal cross-sections, *z*-stacks were acquired with a 1.194 μm *z* distance.

### Image processing and analyses

#### Registration algorithm

Fijiyama registration FIJI plugin (Fernandez and Moisy, 2020), a registration tool for 3D multimodal time-lapse imaging, was used with 4 steps of automatic rigid registration for each alignment of raw samples: 2 steps without, and 2 steps with subpixel accuracy, oversized images subsampling was left on. HuC-channel of all samples was registered to a select HuC signal from a Tg[vglut2a:DsRed;HuC:GFP] 72hpf embryo. Computed transform was then applied to the signals of interest using the “apply a computed transform to another image” option of the Fijiyama plugin.

#### Segmentation algorithm

The Trainable Weka Segmentation FIJI plugin (Arganda-Carreras et al., 2017), a machine learning tool for microscopy pixel classification, was used for image segmentation. Training features for the algorithm were kept to mean and variance in order to limit computing power usage, ensuring accessibility. First, classifiers from the very same embryo were generated from supervised training of the algorithm by different users. Tg[HuC:Kaede] embryos, in which Kaede was photoconverted at 24 hpf, were imaged at 72 hpf. The classifiers generated by the users were then used to segment each sample and to obtain the users’ averaged 3D-model from a single embryo. This process was repeated for each embryo (*n* = 5 embryos per sample). Training took about 4 min per step with 3-4 steps to complete, and generation of a mask through a single classifier took about 10 min on a consumer-grade workstation with an 3700X (8 cores, 16 threads, 3.6Ghz) AMD processor and RAM 32GB.

#### Processing

Users’ and embryos’ averaging, as well as consensus thresholds were performed in FIJI. Homemade batch-processing macros were used to process the whole datasets at once (see Supplementary Material 2). Macros for batch processing of each of the steps were kept independent to ensure easy adaptability, such as variation in sample size or selected thresholds. Signal consensus among all users was used for both quantifications and models.

Signal consensus among 3 or more embryos was used for the generation of the models. Intersection of 3D-models was performed using the image calculator option in FIJI.

#### Tissue volume quantification

Embryos’ 3D-models were registered together and resliced along different axes. Slices were then quantified using FIJI’s 3D object counter. The slicing and quantification of individual slices was automatized with a simple homemade FIJI macro (see Supplementary Material 2). Quantification along axes was processed through R. We used ratios of signal of interest vs. total differentiation domain volumes of individual embryos to generate a nonparametric regression curve with local linear kernel estimation method (Fan, 2018). The R (R core team 2020) package npregfast version 1.5.1 (Sestelo et al., 2017) and the Epanechnikov kernel (with bandwidth parameter selected by cross-validation) were used.

### Photoconversion experiments

HuC:Kaede^Green^ to HuC:Kaede^Red^ photoconversion was carried out at different embryonic stages (24, 36, 48 and 72 hpf) with UV light (λ = 405 nm) using a 20× objective in a Leica SP8 system. Upon exposure to UV light Kaede protein irreversibly shifts emission from green to red fluorescence (516 to 581). Proper photoconversion was monitored by the appearance of strong Kaede^Red^ signal under excitation with a 543 nm laser, and the disappearance of Kaede^Green^. Photoconverted zebrafish embryos were either imaged or returned to embryo medium with phenylthiourea (PTU) in a 28.5°C incubator to let them grow until the desired stage.

### Whole-mount *in situ* hybridization

Embryo whole mount *in situ* hybridization was adapted from (Thisse and Thisse, 2008). The *gad1b* riboprobe encompassing the nucleotides 1104-2334 of accession noAB183390 as described in (Zhang et al., 2020) was used. The *gad1b* template was generated by PCR amplification by adding the T7 promoter sequence (indicated by capital letters) to the reverse primer (*gad1b* Fw 5’— gat ggt tgc gcg gta taa —3’; *gad1b* Rev 5’— ata tta ata cga ctc act ata gCT TCG TTA AAA GGG TGC —3’). The *vglut* probe was as described in (Higashijima et al., 2004). For fluorescent *in situ* hybridization, the DIG-labeled probe was detected with TSA-Cy3. Cell nuclei were stained with DAPI.

## ACKNOWLEDGEMENTS

The authors thank the former and current members of the Pujades lab for help in the training of the algorithms for Trainable Weka Segmentation and for critical insights. We wish to thank Dr S Higashijima (National Institute for Basic Biology, Japan) and Dr. G Sumbre (IBENS) for kindly providing the transgenic lines Tg[gadb1:DsRed] and Tg[vglut2a:DsRed].

## DATA AVAILABILITY

All datasets supporting our work are included as Supplementary Figures and/or are available in our web site https://www.upf.edu/web/pujadeslab

## AUTHOR CONTRIBUTIONS

MB and CP contributed to the conceptualization and design of experiments and analysis of results. MB carried out all the experiments and established the pipeline. FU contributed to the analysis of results. MB and CP wrote the manuscript.

## SUPPLEMENTARY INFORMATION

**Supplementary Figure 1:**
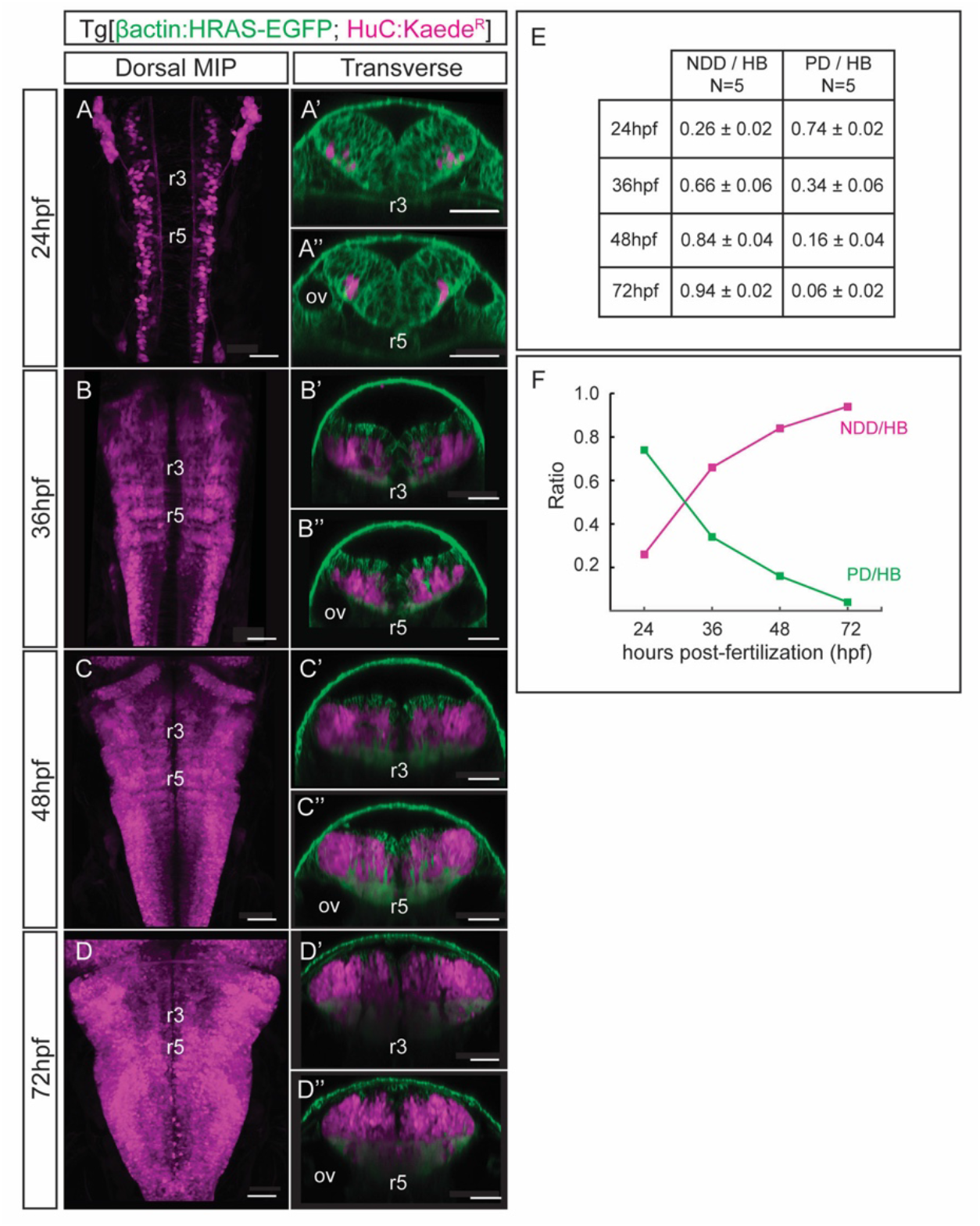
The neuronal differentiation domain growths at the expense of the progenitor domain during hindbrain morphogenesis. (A–D) Dorsal maximal intensity projection (MIP) and transverse views through r3 and r5 of Tg[βactin:HRAS-EGFP;HuC:Kaede] embryos, in which Kaede^Green^ was photoconverted to Kaede^Red^ at the indicated times. Embryos were immediately imaged after photoconversion. Note that the neuronal differentiation territory (NDD in magenta) grows over time, while the progenitor domain (PD in green) decreases. (E) Table representing the mean and SD of volumes’ ratios of the differentiated neurons (NDD/HB) and the progenitor domain (PD/HB) over the total hindbrain, resulting from the quantification of images shown in (A–D); *n* =5 embryos/stage). The quantification of PD was performed by subtracting the HuC-Kaede^Red^ domain to the overall Tg[βactin:HRAS-EGFP] signal. (F) Graph displaying the mean volumes’ ratio.

**Supplementary Material 1: FIJI Macros**

**Supplementary Material 2: R code - Axis distribution plots**

## VIDEOS

**Video 1: 3D-models corresponding to Figure 1.**

(A) Mu4127 3D-model displayed as 360° rotation. Note the volume that r3 and r5 occupy in the whole embryo. (B) Progenitor (nestin in magenta) and neuronal differentiation (HuC in green) domains 3D-models displayed as 360° rotation. The progenitor domains results from the subtraction of the differentiated domain to the nestin signal. (C) neurog1 3D-model displayed as 360° rotation, showing the neurog1-committed progenitors and their derivatives.

**Video 2: 3D-models corresponding to Figure 2.**

(A) Motoneuron (isl1) 3D-model displayed as 360° rotation. (B) Glutamatergic neurons (vglut2a) 3D-model displayed as 360° rotation. (C) GABAergic neurons (gad1b) 3D-model displayed as 360° rotation. (D) Overlay of the three color-coded differentiated neuronal populations 3D-models displayed as 360° rotation.

**Video 3: The temporal registration 3D-atlas**

3D-model of the neuronal differentiation domains contributed by neurons born between 24 and 36hpf (A, green), between 36 and 48hpf (B, blue), and between 48 and 72hpf (C, yellow), displayed as 360° rotation. These 3D-models were obtained by subtracting the model of embryos photoconverted at (A) 24hpf to the model of embryos photoconverted at 36hpf, (B) 36hpf to the model of embryos photoconverted at 48hpf, and (C) 48hpf to the model of the differentiated domain at 72hpf.

**Video 4: The temporal 3D-atlas as a proxy to reveal the birthdate of distinct neuronal differentiated neurons.**

(A) Intersection of the glutamatergic (A) and GABAergic (B) neurons 3D-models and the temporal 3D-map. Intersectional maps are displayed as 360° rotation.

